# A fast numerical method for oxygen supply in tissue with complex blood vessel network

**DOI:** 10.1101/2020.07.02.184259

**Authors:** Yuankai Lu, Dan Hu, Wenjun Ying

## Abstract

Oxygen field evaluation is important in modeling and simulation of many important physiological processes of animals, such as angiogenesis. However, numerical simulation of the oxygen field in animal tissue is usually limited by the unusual coupling of different mechanisms, the nonlinearity of the model, and the complex geometry of refined blood vessel networks. In this work, a fast numerical method is designed for the simulation of oxygen supply in tissue with a large-scale complex vessel network. This method employs an implicit finite-difference scheme to compute the oxygen field. By virtue of an oxygen source distribution technique from vessel center lines to mesh points and a corresponding post-processing technique that eliminate the local numerical error induced by source distribution, square mesh with relatively large mesh sizes can be applied while sufficient numerical accuracy is maintained. The new method has computational complexity which is slightly higher than linear with respect to the number of mesh points and has a convergence order which is slightly lower than second order with respect to the mesh size. As an example, the oxygen field of a tissue irrigated by a blood vessel network with more than four thousand blood vessels can be accurately computed within one minute with our new method. The new method will definitely promote further researches based on evaluation of oxygen field, such as modeling of angiogenesis and pathogenesis of many cardiovascular diseases.

## 1 Introduction

Oxygen plays a key role in animal metabolism. Oxygen supply to tissue is mainly achieved by the circulation system of animals. In particular, the efficiency of oxygen delivery is mainly determined by the microcirculation structure. In order to improve the efficiency of their microcirculation structure, animals have developed different physiological processes to modify the geometry and topology of vessel networks, including blood vessel adaptation and angiogenesis [1, 2]. In these physiological processes, the oxygen level is the key driving stimulus. For example, poor microcirculation structure results in local tissue hypoxia (low concentration of oxygen), which leads to production of growth factors for angiogenesis, such as vascular endothelial growth factor (VEGF) [3, 4]. Angiogenesis plays a critical role in many pathological processes, such as tumor growth [5], wound healing [6], and keloid development [7]

There have been a few techniques for measurement of oxygen level, such as two-photon phosphorescence lifetime microscopy that can be applied to measure oxygen level in *vivo* [8]. However, existing techniques can hardly offer a complete spatial-temporal picture of oxygen field on microscopic scales. Therefore, theoretic modeling [9–11] and numerical simulation [12–17] is widely ultilized in evaluating oxygen level and studying angiogenesis.

The classical Krogh cylinder model roughly describes the oxygen transport from blood vessels to tissues [9, 18]. In this model, evenly spaced capillaries are assumed to be parallel and supply oxygen to a cylindrical tissue domain. Following Krogh’s model, more detailed oxygen consumption mechanisms are taken into account [19]. Coupled models for oxygen delivery including a reasonable oxygen consumption mechanism, a relatively complex vessel network structure, and detailed blood flow in the vessel network are also proposed in later works [10, 20].

Experimental results have indicated disordered spatial distribution of blood vessels [21]. In retinal vascular network, the arterioles and venules reach out from the center to the periphery, forming a roughly radial skeleton, while capillaries form a vessel network that links the arterioles and venules [22]. It is also observed that the microcirculation in tumor tissue presents a chaotic geometry [23]. Therefore, in most real applications, it is important to incorporate the complex vessel network structures into the model and numerical algorithms to evaluate the oxygen field in tissue.

The chaotic vessel geometry brings great challenge in designing effective numerical algorithms. Although existing simulations based on finite difference method and rectangular grids can provide useful insight for tissue oxygen supply, they introduce unrealistic requirement for the geometry of blood vessel networks [2, 13]. A finite element method is also introduced for more general vessel network topology and geometry [19]. This method requires a particular local processing near blood vessels. Meanwhile, the mesh grid is generated according to the vessel geometry. As a result, the corresponding computational cost is huge for tissue with large-scale complex vessel network structure. In more realistic applications, Secomb and Hsu developed a numerical method based on Green’s function, in which each vessel is regarded as a line source of oxygen [17, 23]. Their numerical method can be used to deal with complex vessel geometry. However, their method also suffers from the high computational cost due to the all-to-all interaction between all elements.

In this work, we develop a fast numerical method to evaluate the coupled system for oxygen delivery in tissue, with particular attention paid to the large-scale complex blood vessel network structures. In our model, the blood vessel is also regarded as a line source for oxygen field. A source distribution technique is used so that we can solve the oxygen field on a square mesh. A post-processing technique is employed to remove the numerical error induced by source distribution, which allows us to use a relatively large mesh size for the square mesh, while sufficient numerical accuracy is maintained. Furthermore, an efficient iteration method is used to deal with the nonlinearity of the coupled system. The convergence order of our new method is slightly smaller than second order with respect to the mesh size and the computational cost is slightly larger than linear with respect to the total number of mesh points. Due to the advantages of this new method, we can accurately evaluate the oxygen field of a tissue with more than four thousand blood vessels in about one minute on a standard 3.7GHz personal computer.

## 2 Modeling

The model for oxygen delivery in blood vessels and tissues is well established in previous works [10, 20]. The model mainly includes a partial differential equation (PDE) for the oxygen field, a linear system for blood flow and blood pressure, and a system of ordinary differential equations (ODE) for the blood oxygen.

### 2.1 Oxygen diffusion in tissue

The oxygen-consuming tissue is a mixture of cells, extracellular matrix and extracellular fluid. Oxygen transport in tissue depends mostly on diffusion. The diffusion constant and solubility of oxygen may vary slightly in different tissue. Here we assume a uniform oxygen diffusivity *D* and uniform solubility *α*. Oxygen diffusion in tissue at steady state satisfies the reaction diffusion equation

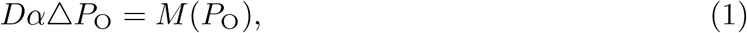

where *P*_O_ is the partial pressure of oxygen and *M* (·) is the oxygen-consuming rate in tissue. The oxygen consumption in cells consists of various biochemical processes, which can be described by the Michaelis-Menten equation in general

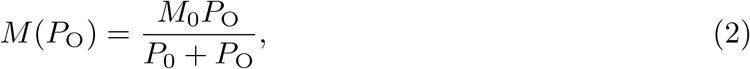

where *M*_0_ represents the maximum consumption under infinite oxygen supply and *P*_0_ represents the partial pressure of oxygen at half-maximal consumption.

### 2.2 Blood flow and blood pressure

The blood flow and blood pressure can be computed with the Ohm’s law and Kirchhoff’s circuit law that form a system of linear equations. For a small blood vessel, which can be regarded as a cylinder, the blood flow in the vessel can be well approximated by the Poiseuille flow. The conductance of the vessel is

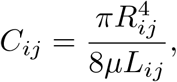

where *i* and *j* are indices of the two end nodes of the vessel, *μ* is the viscosity of the blood, and *R_ij_* and *L_ij_* are the radius and length of the vessel, respectively. The blood pressure and blood flow rate from node *i* to node *j*, *Q_ij_*, satisfies the Ohm’s law

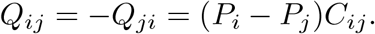

The Kirchhoff’s circuit law describes the conservation of mass

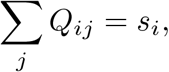

where *s_i_* gives the sources and sinks at the inlets and outlets of the vessel network. At bifurcation nodes or junction nodes, we have *s_i_* = 0. In real applications, the boundary conditions at the inlets and outlets can also be replaced by other conditions, such as fixed pressure condition.

It is worth noting that, the effective viscosity *μ* can be dependent on the vessel radius and blood oxygen level [17, 23]. In this case, the system becomes a nonlinear system that is coupled to whole system for oxygen delivery.

### 2.3 Oxygen flux in vessels

Oxygen carried by blood includes two parts, the minor of which is directly dissolved in the plasma while the major of which is associated with hemoglobin. The relation between oxygen saturation *S_a_* and blood oxygen partial pressure *P*_b_ satisfies the Oxygen-hemoglobin dissociation relation

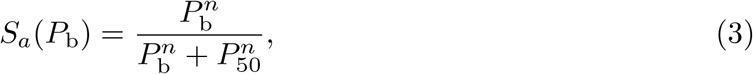

where *n* = 2 is used in this work and *P*_50_ is the half-saturated oxygen pressure that may depend on the pH-value of blood. Correspondingly, the oxygen flux *f* through a cross-section of the vessel includes two parts

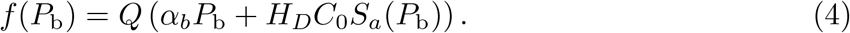

where *Q* is the blood flow rate, *α_b_* is the oxygen solubility in blood plasma, *H_D_* is the discharge hematocrit, and *C*_0_ is the concentration of hemoglobin-bound oxygen in a fully saturated red blood cell (RBC).

### 2.4 Oxygen exchange on vessel walls

Let *s* be the arc-length parameter along the center line of a vessel and **x**(*s*) be the coordinate of the center line. Assume the cross section of the blood vessel is a circle with radius *R*, then the total oxygen flux *q*(*s*) through the blood vessel wall per unit length is

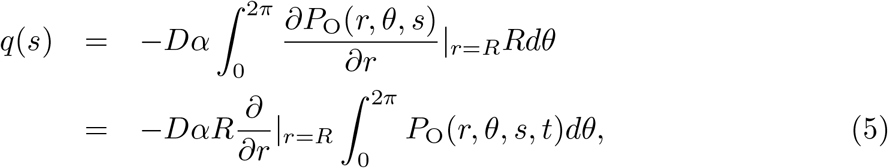

where *r* is the polar radius and *θ* is the polar angle on the cross-sectional plane (the pole is at **x**(*s*)). Ignoring the oxygen consumption in blood, we obtain the conservation of oxygen

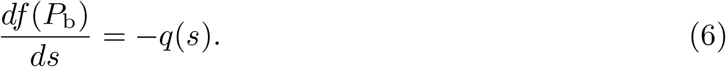

Given the oxygen flux *q*(*s*), the above equation becomes a first order ordinary differential equation (ODE). A boundary condition is required to solve the equation for each vessel. In real applications, we may have *P*_b_ at the inlet of each vessel: (1) At the inlets of the vessel system, *P*_b_ is given; (2) At the bifurcation points, *P*_b_ at the inlet of the downstream vessels is inherited from the parent vessel; (3) At collecting junctions, *P*_b_ at the inlet of the downstream vessel is obtained from its parent vessels as the mixed value by conservation of oxygen.

### 2.5 Model simplification

Since the oxygen partial pressure should be continuous in the whole domain, i.e., the tissue domain and the vessel domain, *P*_b_ provides a Dirichlet boundary condition on vessel walls for *P*_O_ in Eq. (1). However, the disordered structure of blood vessel walls brings great difficulties in meshing and numerical simulations to solve the above coupled system. In previous studies [24], the oxygen flux are represented by oxygen sources on the center lines of vessels. Under this simplification, the governing equation for oxygen supply is defined in the whole domain. In this case, the governing equation becomes

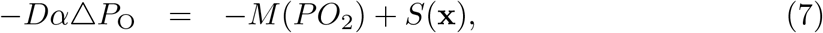

where *S*(**x**) = ∫ *q*(*s*)*δ*(**x** − **x**(*s*))*ds* is the oxygen source supplied by the vessel. Note that the vessel radius is relatively small (2 ~ 3*μm*) compared to the distance between capillaries (~ 100*μm*). Therefore, this simplification can be a good approximation at least for far field. The oxygen partial pressure may be overestimated near the vessels, which can lead to a slight overestimate of the oxygen source *M* (*P*_O_). Further improvement on such an approximation has been discussed in the work of Ref. [17, 25].

Now we have obtained a coupled system for oxygen delivery, which includes a PDE (7) on the whole domain, a set of nonlinear ODEs (6) on the vessel centerlines, and a system of linear equations for blood pressures and blood flows in all vessels.

## 3 Numerical method

In order to develop a fast numerical method to solve the above coupled system, we are left with two main tasks: (1) find an efficient iteration method to deal with the nonlinearity of the coupled system and (2) develop a fast numerical solver for the PDE (7) with complex sources on blood vessel centerlines.

### 3.1 Nonlinear iteration

Mainly due to the nonlinearity of the ODEs (6) and the unusual form of coupling, general iteration methods, such as the full Newton-like iterative methods, are both hard in code implementation and lack of convergence guarantee for the coupled system. Here we propose an iterative method by decoupling the system and alternatively updating *P*_O_ and *P*_b_. In particular, we introduce pseudo time step in order to reach the steady state solution. The rest of the section then explain how each (pseudo) time step is solved, namely with a semi-implicit Euler method.

Given 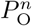 and 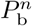 at step *n*, by assuming that the blood oxygen partial pressure *P*_O_ = *P*_b_ on vessel walls, Eq. (5) allows us to evaluate the flux *q^n^*(*s*) numerically. In the 3-D case, we may compute the integral in Eq. (5) first and then use a linear fitting along the *r*–direction to evaluate the derivative in Eq. (5). In the 2-D case, linear fitting on both sides of the vessel can be directly utilized to evaluate the flux *q^n^*(*s*). Detailed discussion on the fitting in the 2-D case is included in Appendix A. In general, we denote the evaluation of *q^n^* as

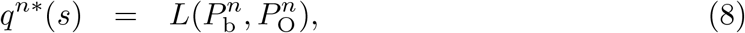

which is linearly dependent on 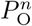 and 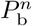. According to our numerical tests, it is not efficient for convergence to update the flux directly using 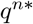. Instead, a weighted sum

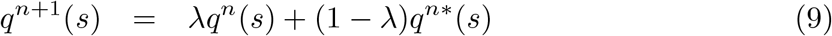

is more efficient, where the weight 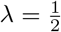 is used in our simulation. Once the flux is obtained, Eq. (6) is used to update *P*_b_ in all vessels

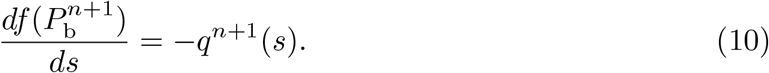

The fourth-order Runge-Kutta method is used to solve the ODE (10) to obtain 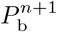 in each vessel segment. In practice, 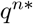 can also be updated in a Gauss-Seidel fashion. Namely, the values of 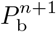 on the updated nodes can be used in Eq. (8) to evaluate the flux.

The flux *q^n^*^+1^ is also used to calculate the oxygen source *S^n^*^+1^, which is required in computing 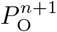. It is possible to update *P*_O_ by fully solving the PDE (7) with given oxygen source *S^n^*^+1^. However, since we still need the nonlinear iteration, it is neither necessary nor efficient to find the accurate solution at each iteration step. Instead, we use the following scheme to update *P*_O_

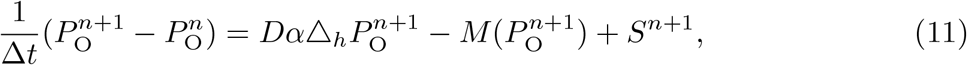

where Δ_*h*_ is the numerical Laplacian operator and Δ*t* is the temporal step size. This implicit numerical scheme solves the time-dependent reaction-diffusion equation for one time step. When the iteration converges, the solution satisfies the steady state PDE (7).

Note that an increase of the oxygen partial pressure 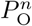 in tissue can lead to a decrease of the oxygen flux *q^n^*^+1^ (thus the oxygen source *S^n^*^+1^). This implies at least one negative eigenvalue of the coupled system in the sense of linearization. As a result, it may even be not stable to update *P*_O_ by fully solving the steady state PDE (7) for each iteration step (namely, Δ*t* = ∞). The instability has been observed in our numerical test. From this point of view, a suitable time step size Δ*t* should be selected so that it is both good for iteration stability (Δ*t* is not too big) and good for fast convergence (Δ*t* is sufficiently big).

### 3.2 PDE solver

Based on the iteration framework above, our remaining task is to numerically solve the PDE (7) efficiently. For the simplified coupled system, we do not consider the detailed geometry of vessel walls. Thus,a square mesh can be easily used in our simulation. We simply use the central difference as the numerical Laplacian Δ_*h*_, where *h* is the mesh size. This brings great convenience in developing a fast solver.

There are two problems left for us to solve: First, the oxygen sources are located on vessel centerlines, we need to distribute the sources onto the square mesh points; Second, in order to conduct a large scale simulation, we need to use a relatively large spatial mesh size while maintaining sufficient numerical accuracy.

In order to distribute the oxygen sources onto the square mesh points, first we discretize the sources to be point sources 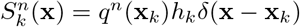 on the center lines, where *k* is the index of the point source at **x**_*k*_ = (*x_k,_*_1_, *x_k,_*_2_, …, *x_k,d_*) (*d* is the space dimension) and *h_k_* is the step size on the centerline; Then, we use Peskin’s numerical *δ*–function 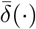 (see Appendix B) to distribute all point sources onto their neighboring mesh points [26]. Namely, we have

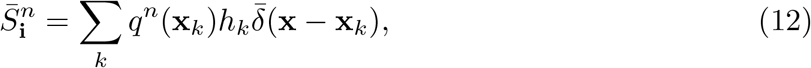

where 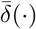 is the numerical *δ*–function and **i** = (*i*_1_*, i*_2_*, …, i_d_*) is the index of the mesh point.

Due to the nonlinearity in the consumption function *M* (·), we use the standard multigrid algorithm combined with the Newton’s method to solve Eq. (11). According to our numerical tests, only a few steps of Newton-iteration is sufficient to make the numerical error small enough.

Notably, the above redistribution of oxygen-sources can lead to numerical error in the solution. As a result of using Peskin’s numerical *δ*–function, the numerical error induced by source redistribution is local — mainly on the local mesh points to which the oxygen sources are distributed (see Appendix C). Nevertheless, the local oxygen field around the blood vessels must be sufficiently accurate for evaluating the oxygen flux using Eq. (8). Therefore, without further improvement, we can only use a small spatial mesh size *h* to perform simulations.

Next, we introduce a post-processing technique to reduce the local error induced by the redistribution of oxygen-sources. With this post-processing step, we are able to use a relatively large mesh size *h* (e.g., be comparable to or even larger than vessel diameters) while maintaining a sufficiently small numerical error.

### 3.3 Post-processing

The post-processing is designed to reduce the error introduced by oxygen-source redistribution. This error can be defined as the difference 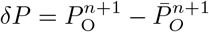, where 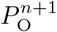 and 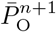 satisfy the following equations

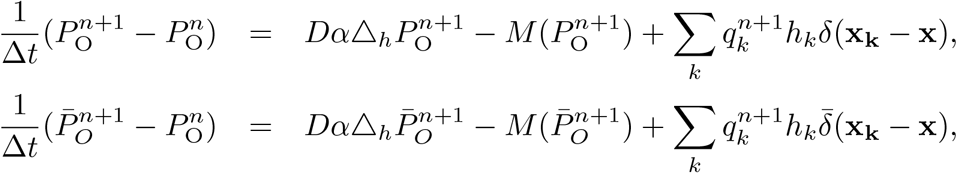

respectively. Therefore, *δP* satisfies

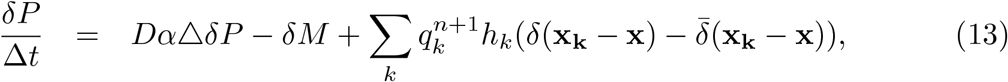

where 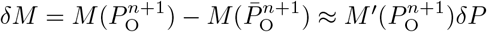.

Equation (13) appears to be nonlinear due to the nonlinearity in *δM*. However, in real applications, this term is always very small and we can neglect it. The reason that *δM* is small is double fold: For mesh points near a vessel, the oxygen partial pressure is relatively high (much bigger than *P*_0_), thus 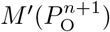 is very small; whereas for mesh points away from all vessels, *δP* becomes very small thanks to the good property of the numerical *δ* function. Therefore, we can evaluate *δP* by neglecting the nonlinear term

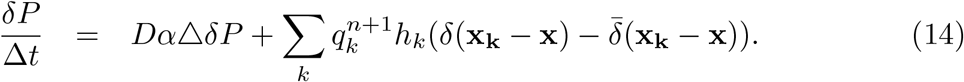

Due to the linearity of Equation (14), *δP* can be regarded as the linear combination 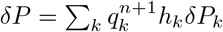, where *δP_k_* satisfies

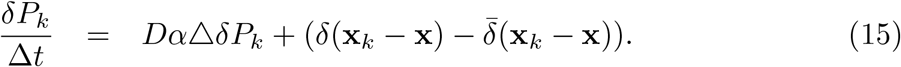

The boundary condition for Eq. (15) is that *δP_k_* vanishes when **x** → ∞. In fact, as shown in Appendix C, only a few mesh-sizes away from **x**_*k*_, the error *δP_k_* is already negligible. This observation allows us to solve *δP_k_* only on local mesh near to **x_k_** (e.g., an 8 × 8 mesh in the 2-D case).

The error *δP_k_* can be further decomposed into two parts 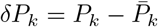, where the first part 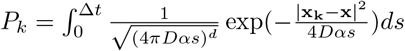 is the fundamental solution corresponding to the point source *δ*(**x_k_** − **x**) and the second part 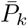 is corresponding to the distributed sources

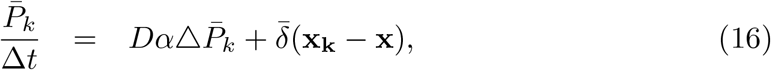

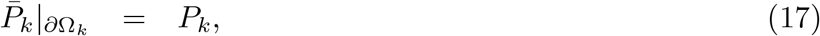

where *∂*Ω_*k*_ is the boundary of the local domain Ω_*k*_. Note that the coefficient matrix to numerically solve the linear system (16) on the local mesh is independent of *k* and the size of the coefficient matrix is small (e.g., 64 × 64). We can find and save the inverse of the coefficient matrix numerically. Then a matrix-vector multiplication is used to find 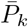.

In summary, for each source point **x_k_**, *δP_k_* is evaluated on a local mesh at the beginning. At each iteration step, the linear combination 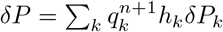 is used to evaluate the error on mesh points close to vessels introduced by oxygen-source redistribution. This error is then removed from 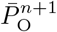.

## 4 Parameters for simulation

The model parameters used in this work are listed in Table 1. The data are adapted from previous experiments on different animal retinas [27–31]. Our numerical method can also be applied to systems with non-uniform parameters.

**Table 1.** Model parameters

| Blood oxygen parameters |  |  |
| --- | --- | --- |
| Maximal RBC oxygen concentration | $C_0 = 0.5 \text{ cm}^3 \text{ O}_2 / \text{cm}^3$ | [17] |
| Effective oxygen solubility | $\alpha_b = 3.1 \times 10^{-5} \text{ cm}^3 \text{ O}_2 / \text{cm}^3 \text{ mmHg}$ | [17] |
| Hill equation parameter | $P_{50} = 38 \text{ mmHg}$ | [17] |
| Hill equation parameter | $n = 2$ | [17] |
| Tissue oxygen parameters |  |  |
| Diffusion constant | $D\alpha = 6 \times 10^{-10} \text{ cm}^3 \text{ O}_2 / \text{cm} \cdot \text{s} \cdot \text{mmHg}$ | [17] |
| Consumption parameter | $M_0 = 1.7 \times 10^{-4} \text{ cm}^3 \text{ O}_2 / \text{cm}^3 \cdot \text{s}$ | [17, 27] |
| Consumption parameter | $P_0 = 1 \text{ mmHg}$ | [17] |
| Blood flow parameters |  |  |
| Oxygen partial pressure at inlet | $P_{b0} = 100 \text{ mmHg}$ | [28] |
| Total flow | $Q_0 = 2000 \mu\text{m}^3 / \text{s}$ | [30] |
| Blood viscosity | $\mu = 4.6 \times 10^{-9} \text{ g} / \text{cm}^2$ | [29] |

## 5 Numerical Results

### 5.1 Blood vessel network structures

Two model structures of blood vessel networks are used in our simulation (see Fig 1). The simple cobweb structure (Fig 1(a)) is used for numerical convergence analysis. The refined structure with 4306 blood vessels (Fig 1(b)) is adapted from real experimental measurements on a mouse retina. It is used for analyzing the efficiency of oxygen supply.

**Fig 1.**
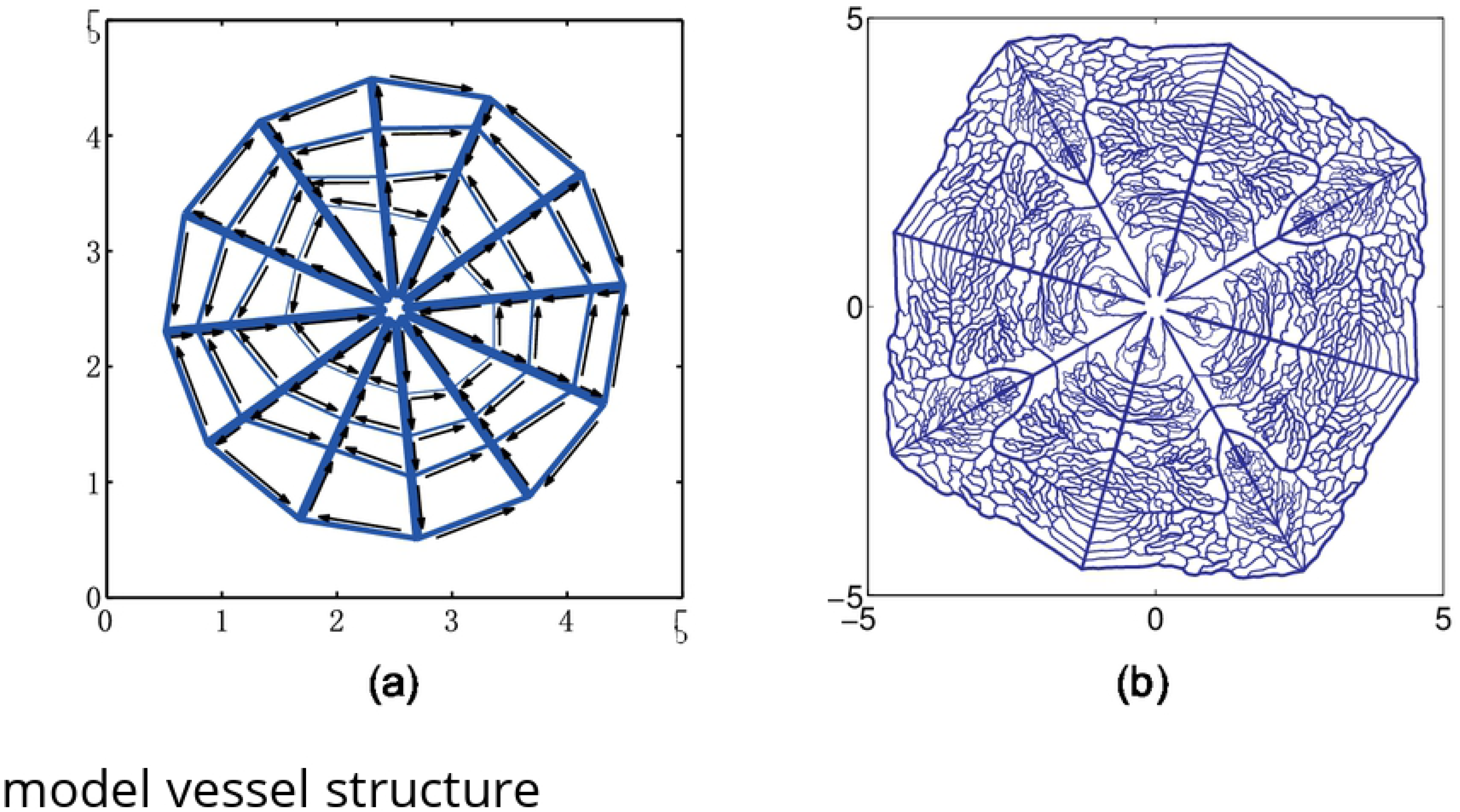
Model vessel structures. (a) A cobweb vessel structure mimicking the main branches of retinal vessel networks. There are six inlets and six outlets for blood flow near the center of the network. The inlets and outlets are in a spaced arrangement. The arrows show the direction of blood flow. (b) A refined vessel network with 4306 blood vessels. The network structure is adapted from a real retinal vessel network measured in experiment by stretching and symmetrical extensions. There are four flow inlets and flow outlets near the center. The widths of the lines in both figure show the radii of blood vessels. The unit for the axes is millimeter.

### 5.2 Numerical solution for the cobweb vessel network

In Fig 2(a), we show the numerical solution of the oxygen partial pressure obtained with a 1024 × 1024 square mesh for the cobweb vessel structure. Since the distance between neighboring vessels is much bigger than that of real micro vasculature, tissue away from the vessels is in a hypoxic state, i.e., the oxygen partial pressure is very low. The width of the well irrigated region for each vessel depends on the blood flow in it, being roughly on the scale of 100 *μm*.

**Fig 2.**
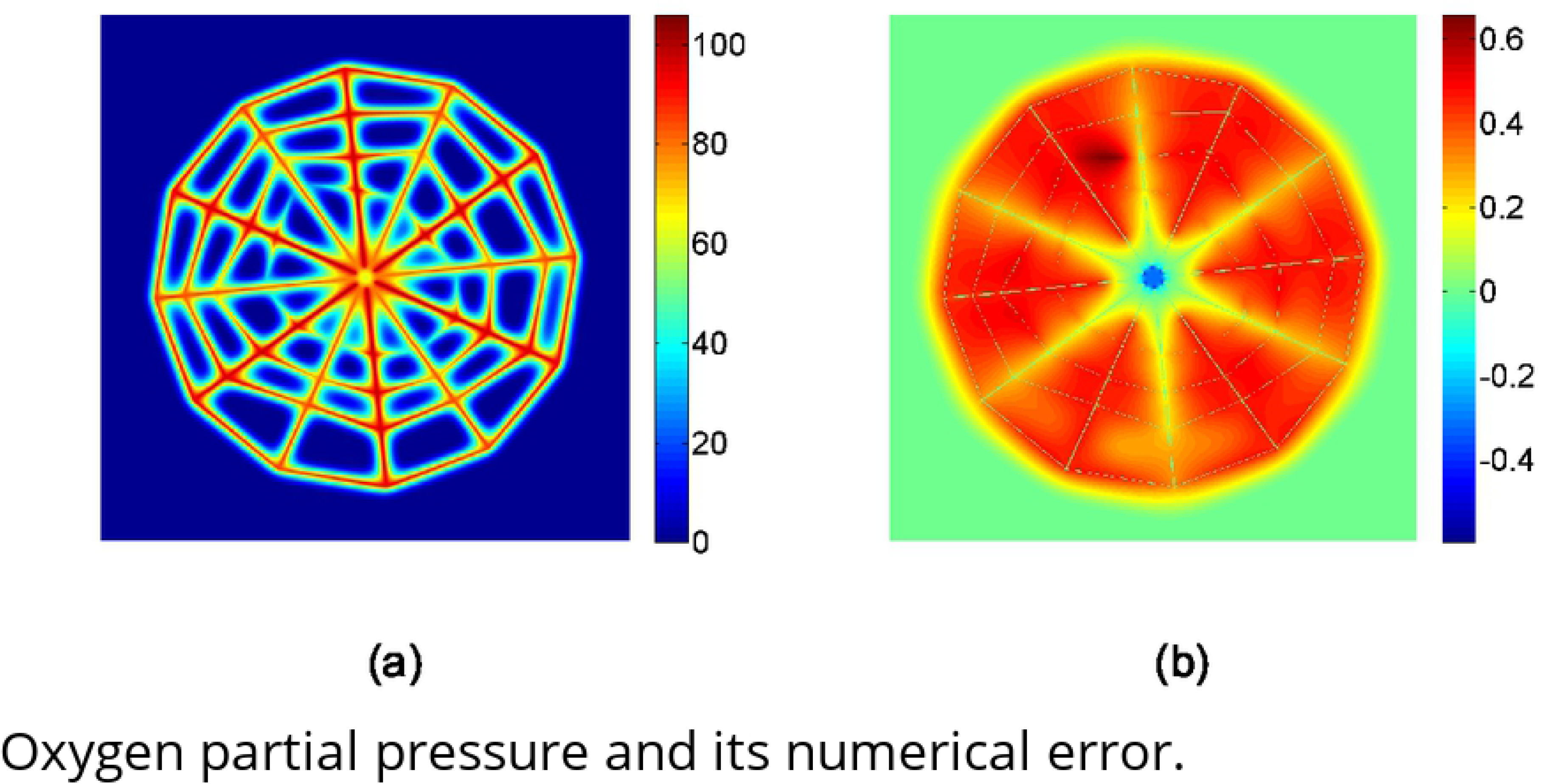
Oxygen partial pressure and its numerical error. (a) Numerical solution of the oxygen partial pressure obtained with a 1024 × 1024 square mesh. (b) Numerical error of the oxygen partial pressure calculated by the difference between the solution obtained with a 1024 × 1024 mesh and a 2048 × 2048 mesh. The unit for oxygen partial pressure is *mmHg*.

The numerical error of the oxygen partial pressure is shown in Fig 2(b). The error is calculated by comparing the solution obtained with a 1024 × 1024 mesh and a 2048 × 2048 mesh. The error is smaller than 1 *mmHg*. This suggests that with a mesh size of about 5 *μm*, the numerical error can be smaller than one percent, which is much smaller than the modeling error in general.

In Fig 3, we show the blood oxygen partial pressure *P*_b_ and oxygen flux *q* on all vessels. Because the blood vessels of the most inner loop are very small, they have a big resistance to blood flow. As a result, the blood flow in these vessels is very small, which leads to a fast decay in blood oxygen partial pressure along the flow direction (see Fig 3(a)). From Fig 3(c), we can see that the blood oxygen partial pressure *P*_b_ decreases along each vessel segment as the oxygen diffuses from vessel to tissue. Similarly, in Fig 3(b) and 3(d), we show the oxygen flux *q*. From Fig 3(b), we can see that the flux *q* reaches minimums at most cross-links of the vessel segments. This is because that more than one vessels “share” the oxygen supply to the local tissue at the cross-links. On the contrary, from Fig 3(d), we can see that the oxygen flux in vessel *a* and *d* reaches the maximums at their one or two ends. This is due to the vacancy of vessels at the center and the outer domain. At the end of the vessel segment *b*, the blood oxygen partial pressure becomes a constant and the flux becomes zero, because the oxygen partial pressure is higher in the surrounding tissue than the blood inside the vessel. Here we set the negative flux to be zero. This setting does not significantly change the tissue oxygen level and blood oxygen concentration in downstream vessels since this piece of vessel is always short.

**Fig 3.**
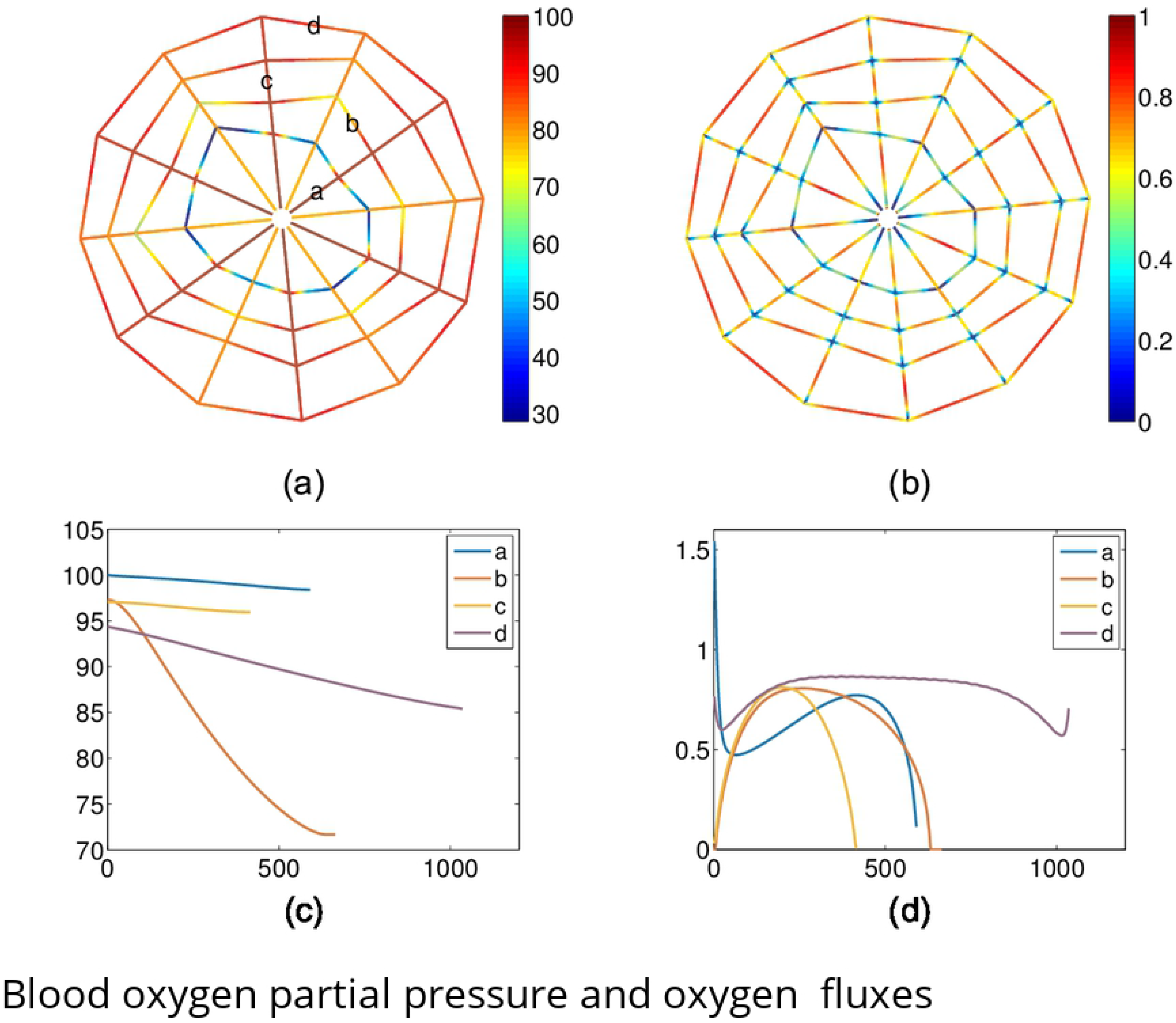
Blood oxygen partial pressure and oxygen fluxes from the vessels. (a) Blood oxygen partial pressure *P*_b_ on the vessels. (b) Oxygen fluxes *q* from the vessels to tissue. (c-d) *P*_b_ and *q* on four marked vessel segments in (a). The *x*–axis is the length from the inlet for vessels *a* to *d*.

### 5.3 Numerical solution for the refined vessel network

The numerical solutions of the oxygen partial pressure *P*_O_ obtained with the refined vessel network with a 1024 × 1024 mesh are shown in Fig 4. The oxygen partial pressure field (a) is obtained with a normal tissue consumption (*M*_0_ = 1.7 × 10^−4^*cm*^3^*O*_2_*/cm*^3^ · *s*), whereas the partial pressure field (c) is obtained with a reduced tissue consumption (*M*_0_ = 2.89 10^−5^*cm*^3^*O*_2_*/cm*^3^ · *s*). In both cases, we can see that the oxygen partial pressure directly reflects the refined vessel structure. In Fig 4 (b) and (d), we statistically analyze the oxygen fields inside the circles shown in Fig 4 (a) and (c), respectively. For the normal consumption case as shown in Fig 4 (a) and (b), tissue in the outer domain has a very low oxygen supply. This is due to the large stretch in the outer domain when we generate the refined vessel network, which significantly affects the oxygen supply in two folds: First, it increases the irrigated area of the corresponding blood vessels in the outer domain; Second, it increases the resistance of these vessels, thus decreases the blood flow in these vessels. For the reduced consumption case as shown in Fig 4 (c) and (d), all tissue inside the circle is well irrigated.

**Fig 4.**
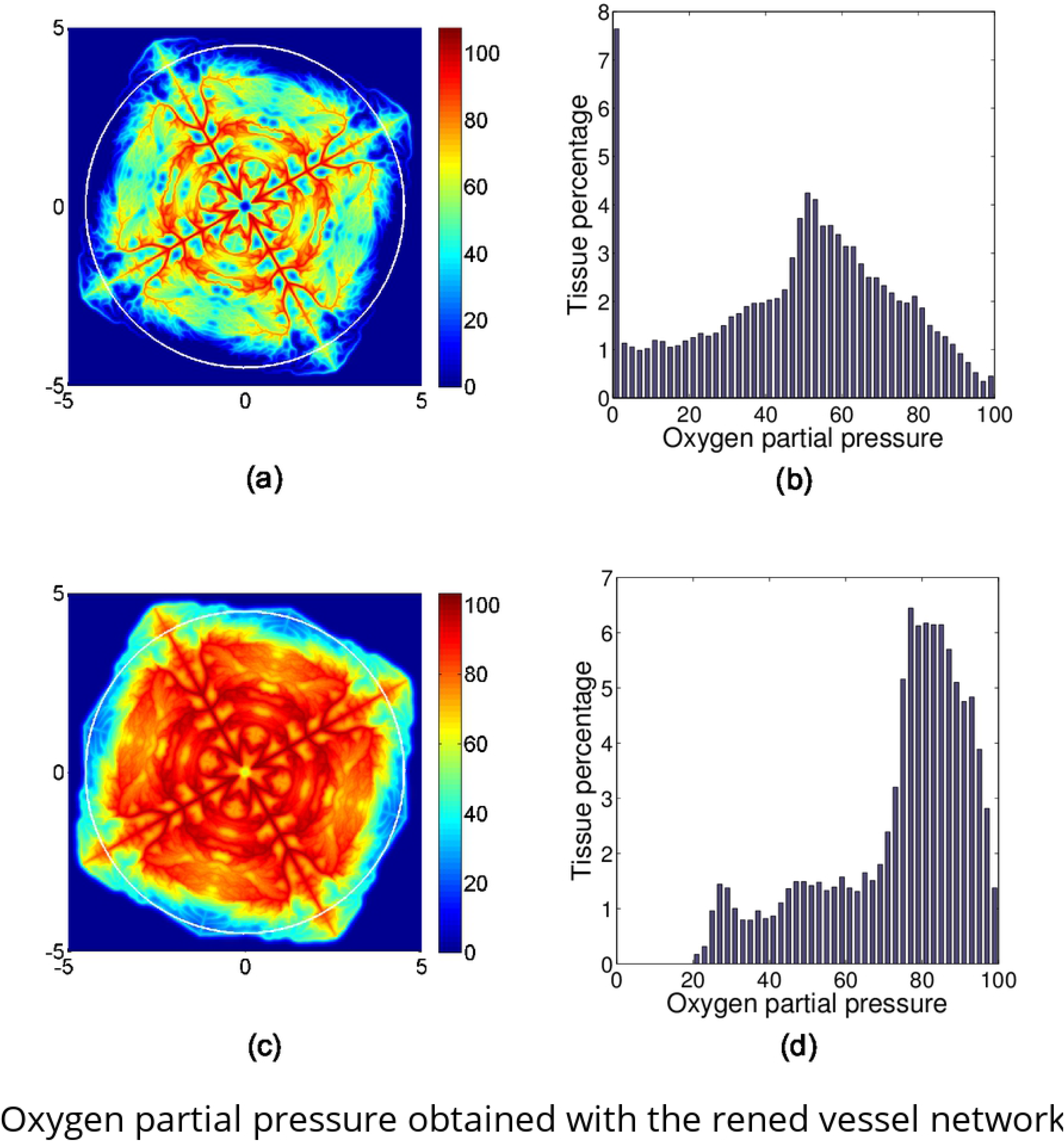
Oxygen partial pressure obtained with the refined vessel network. The partial pressure for normal tissue consumption and reduced tissue consumption are shown in (a) and (c), respectively. The tissue inside the white circles shown in (a) and (c) are used to statistically evaluate the area percentages of tissue with particular partial pressure of oxygen. The statistical results are shown in Figure (b) and (c), respectively.

### 5.4 Convergence analysis and efficiency analysis

The simple cobweb vessel network is used to numerically study the convergence of our new method. The results for mesh refinement test are shown in Fig 5(a). The relative error *E_n_* is computed by the normalized *L*^2^–norm of the difference between the solutions obtained with an *N × N* mesh and a 2*N* × 2*N* mesh. The numerical convergence order is about 1.73 with respect to the mesh size. When *N* = 1024, the relative error is smaller than 1%, which is consistent with the result shown in Fig 2(b). The iteration convergence is shown in Fig 5(b), where the relative difference is shown by the normalized *L*^2^–norm of the difference of the corresponding functions, *q* and *P*_O_, between two iteration steps. We can see that the relative differences have a fast decay at the beginning, followed by a relatively slower linear convergence. It is sufficient to make the relative difference smaller than 1 × 10^−3^ with less than 50 iteration steps.

**Fig 5.**
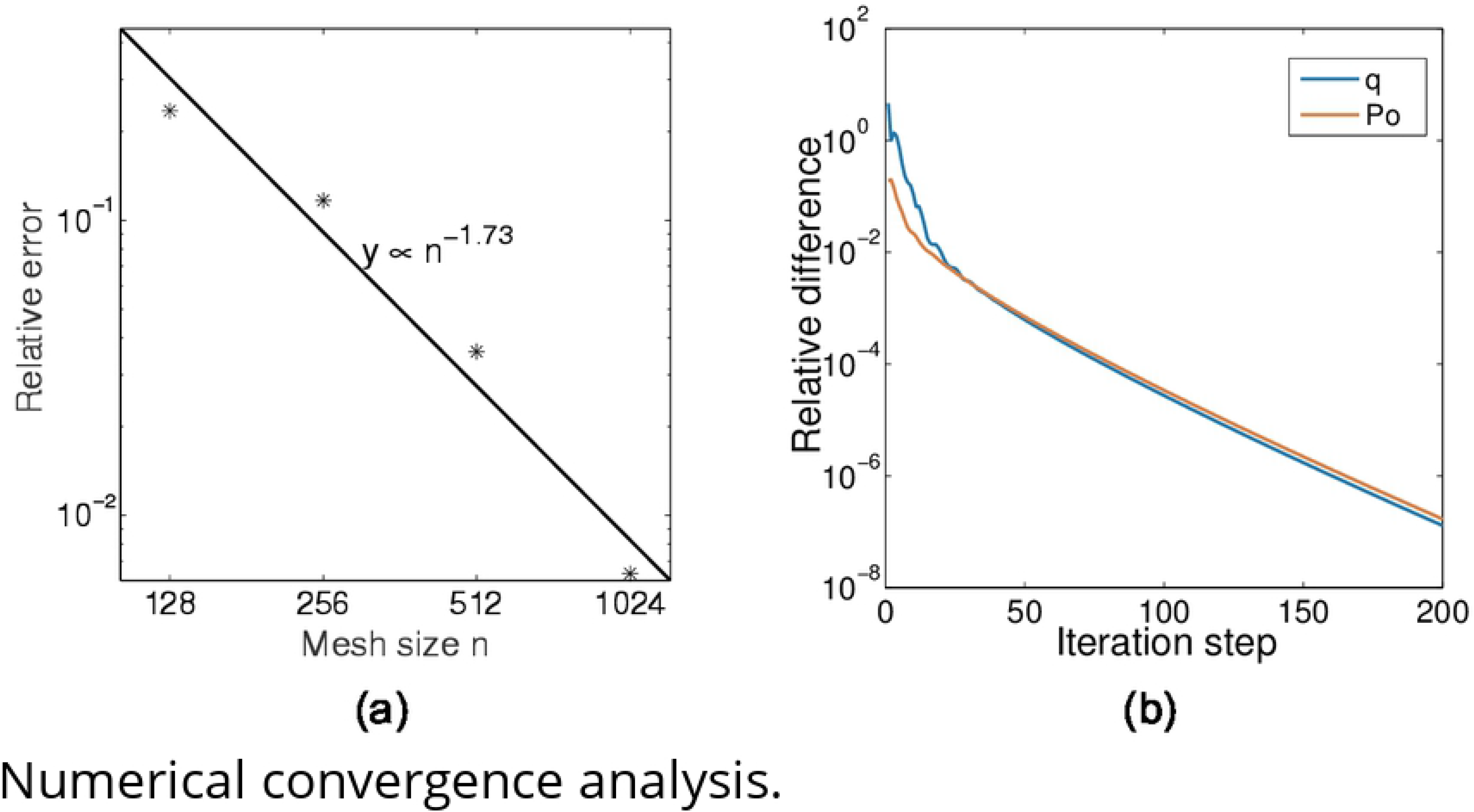
Numerical convergence analysis. (a) Mesh refinement test. The stars show the numerical error. The fitted line shows a convergence order of 1.73 with respect to mesh size. (b) Iteration convergence.

The time cost is analyzed in Fig 6. At the beginning of each iteration, three to six Newton iterations are required for each full iteration step. In each Newton iteration step, two to three V-cycles of multigrid iterations are required to solve the oxygen partial pressure field. After a few full iteration steps, only two to three Newton iteration steps, each with only one V-cycle, are sufficient to make the error small enough for each iteration step. In Fig 6(a), we show the average time cost for each Newton iteration step, tested on a standard 3.7 GHz personal computer. Note that the discrete step sizes *h_k_* along the blood vessels are proportional to the mesh size. Hence the total number of discrete nodes on the blood vessels is proportional to *N*, while the total number of spatial nodes for the oxygen field is proportional to *N* ^2^. As a result, the average time cost in solving the PDE for *P*_O_ increases much faster than that in solving the ODEs for *P*_b_ and the post-processing. Even for the refined vessel network, the time cost in solving the ODE and post-processing is only a small part in the total time cost for large *N* (e.g., 1024).

**Fig 6.**
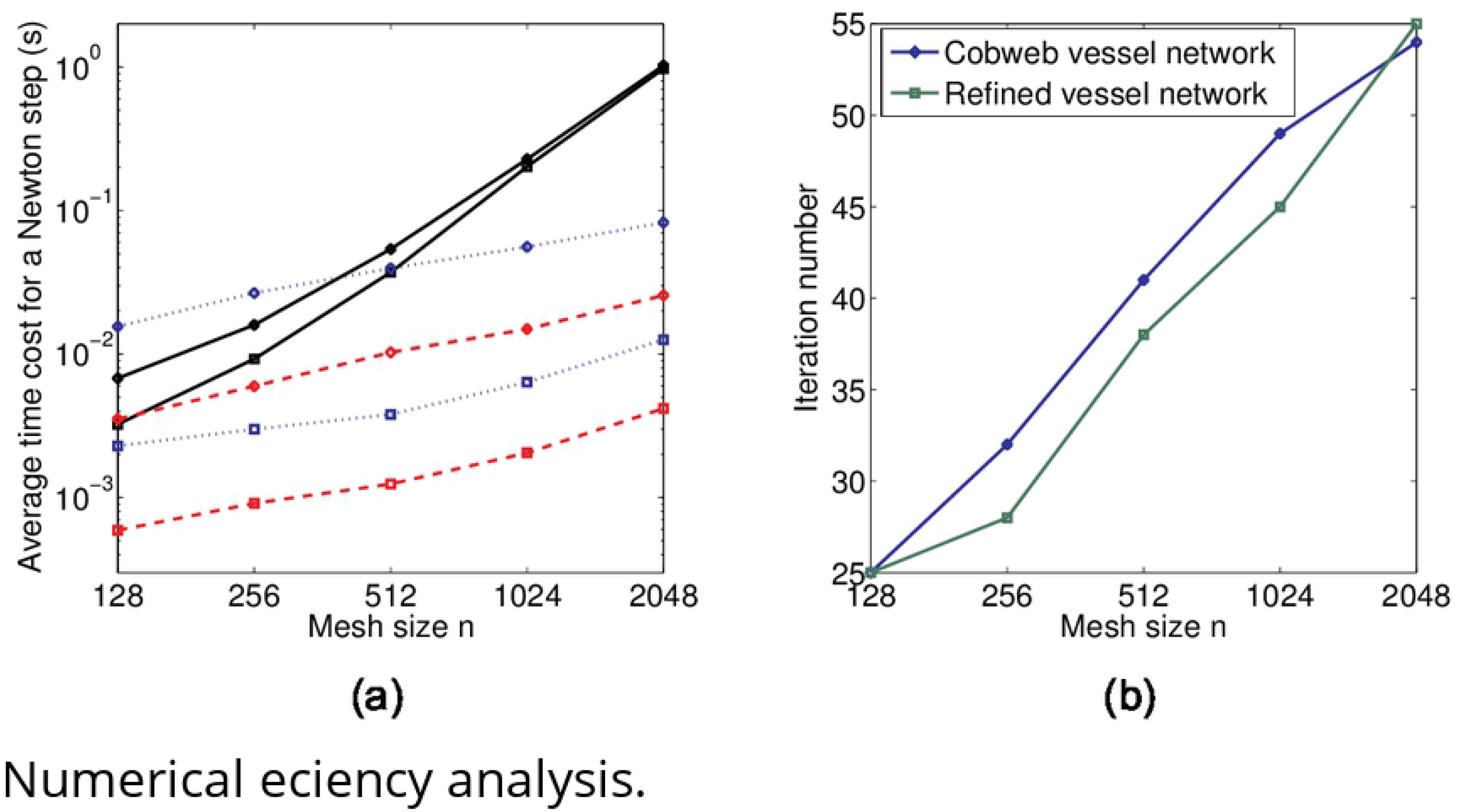
Numerical efficiency analysis. (a) Average time cost for each Newton iteration step. The black solid lines, the blue dotted lines, and the red dashed lines show the average time cost for the PDE, the ODEs, and the post-processing in each Newton iteration step, respectively. The circled and squared lines show the time cost for the refined vessel network and the simple cobweb network, respectively. (b) Iteration numbers for different mesh-size.

The total number of full iterations used to make the relative difference smaller than 1 × 10^−3^ is shown in Fig 6(b). We can see that the iteration numbers is comparable for the simple and refined vessel network structure, while both of them increase slightly with the mesh size *N*. The increase in iteration numbers is mainly due the choice of the temporal step size Δ*t* in Eq. (11). As we have discussed above, a large Δ*t* is helpful for convergence while a too large Δ*t* can induce instability in iteration. According to our numerical test, the optimal Δ*t* decreases with *N*. For example, Δ*t* ≈ 5000 is used for *N* = 1024 in our simulation, while Δ*t* ≈ 3500 is used for *N* = 2048.

The average number of Newton iterations required for each full iteration step is about 5 ∼ 6. Therefore, the time cost is about 40 seconds to find the solution with a relative error smaller than 1% on a 1024 × 1024 mesh with a standard 3.7 GHz personal computer

## 6 Conclusions and Discussions

Oxygen delivery in tissue plays an important role in many physiological processes such as angiogenesis, blood flow regulation, and blood vessel adaptation. Repeatedly evaluating the oxygen field in tissue is the key bottle-neck that limits the large scale modeling and simulation of these important processes. In this work, a fast numerical method is developed for the computation of the nonlinear coupled system of oxygen consumption, oxygen diffusion, and oxygen delivery in blood vessels with complex network structures.

Our fast numerical method involves an implicit finite-difference method for solving the partial differential equation of oxygen partial pressure with a square mesh. The key techniques we used includes (1) the Peskin’s numerical *δ*–function to distribute the oxygen sources onto mesh points and (2) the post-processing to remove the numerical error induced by the distribution of oxygen sources. With these techniques, relatively large spacial mesh size can be used while sufficient numerical accuracy is maintained. The computational complexity is slightly bigger than linear with respect to the number of mesh points, taking into account the increase in iteration steps for refined meshes. The convergence of numerical error is slightly less then second order with respect to the mesh size. With a 1024 × 1024 square mesh, the numerical error is smaller than 1%, which is typically smaller than the modeling error in applications. The time cost to obtain the numerical solution is about 40 seconds on a 1024 × 1024 mesh and about 170 seconds on a 2048 × 2048 mesh on a standard 3.7 GHz personal computer. Our numerical method can also be naturally extended to three-dimensional simulations.

Nevertheless, we can see that the total number of iterations used in our current simulations is still large. Hence, to pave the way for more realistic three dimensional simulations with complex blood vessel networks, better iteration strategies should be explored to reduce the time cost in our method.

## 7 Acknowledgement

This work is supported by the National Natural Science Foundation of China (Contract no. 11971312, 11771290, and 91630208) and Student Innovation Center, Shanghai Jiao Tong University.

## A Oxygen flux through blood vessel walls

For the two-dimensional case, oxygen flux is evaluated on both side of the vessel walls separately. The oxygen partial pressure *P*_O_ is assumed to be equal to *P*_b_ on vessel walls. In order to evaluate the oxygen flux at a discrete node **x** = (*x, y*) on the vessel center line, we first numerically calculate the unit normal direction **n** by central difference. Then, we define two corresponding points **x**^±^ = **x** ± *r***n** on the vessel wall (see Fig 7), where *r* is the vessel radius. Next, the oxygen partial pressures *P*_O_ on four nearest mesh points to **x**^+^ outside the vessel are used to fit a linear function *P*_O_(*x, y*) ≈ *P*_b_ + **G** · (**x** − **x**^+^) in the sense of least squares, where **G** is the gradient to be fitted. Finally, the oxygen flux is given by *q* = *Dα***G** · **n**.

**Fig 7.**
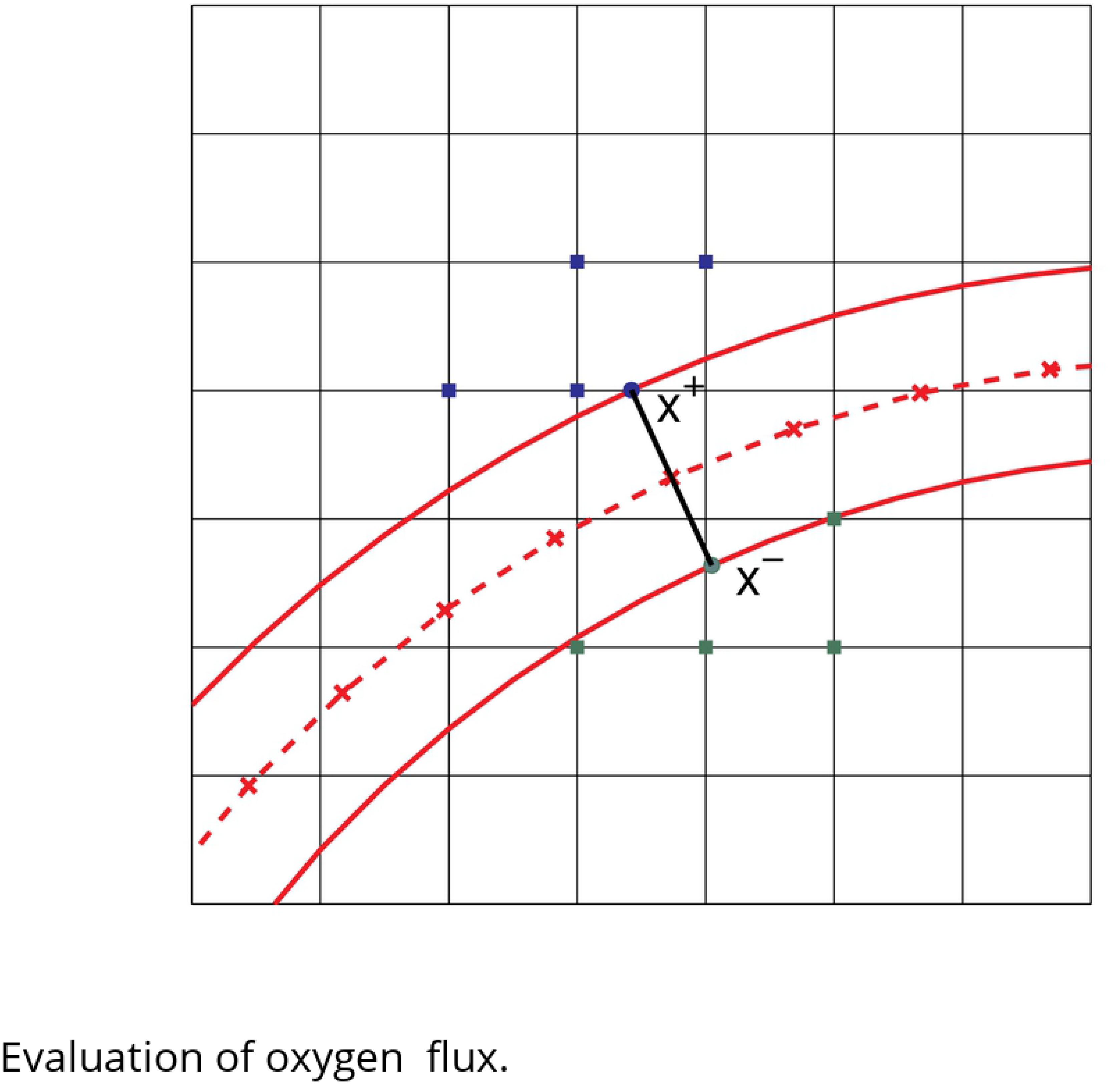
Evaluation of oxygen flux. The red solid lines represent the vessel wall. The red dashed line shows the center line of the blood vessel, the crosses on which show the discrete nodes. The blue (green) solid circle shows *x*^+^ (*x*^−^), whereas the blue (green) solid squares show the nearest mesh points used to fit the linear function and evaluate the flux.

## B Peskin’s numerical *δ*−function

Peskin’s numerical *δ*−function is defined as

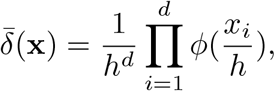

where *d* is the dimension of **x**, h is the mesh size, and *x_i_* are the Cartesian components of **x**. The one-dimensional continuous function *ϕ*(*x*) is given by

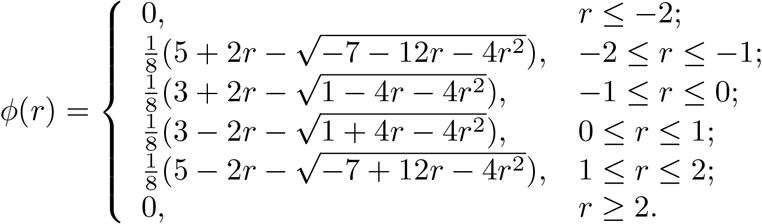

*ϕ*(*x*) satisfies the following constraints

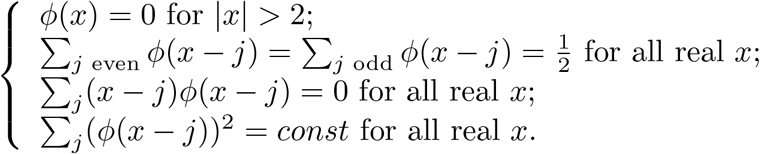

## C Localization of *δP_k_*

An example of *δP_k_* defined in Eq. (15) on the local mesh is illustrated in Fig 8, where the mesh size is *h* = 0.002, **x_k_**= (0.5 + *h/*2, 0.5 + *h/*2) is set at the center of the local mesh. It can be clearly seen that *δP_k_* decays very rapidly. When |**x** − **x_k_**| > 2*h*, the error is already negligible.

**Fig 8.**
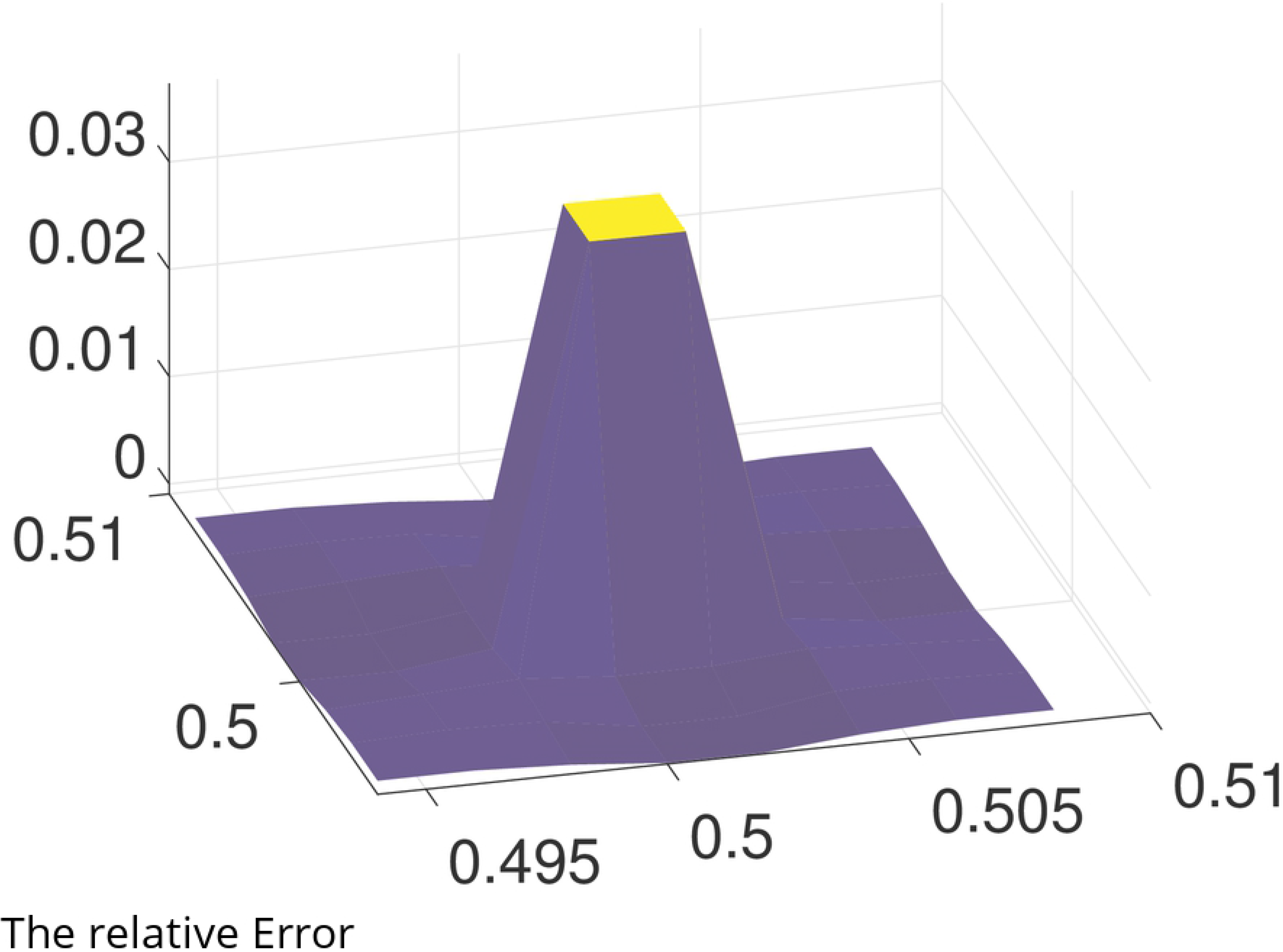
The relative Error *δP_k_* on a local mesh.

## References

1. Secomb TW, Alberding JP, Hsu R, Dewhirst MW, Pries AR. Angiogenesis: an adaptive dynamic biological patterning problem. PLoS computational biology. 2013;9(3):e1002983.

2. Bartha K, Rieger H. Vascular network remodeling via vessel cooption, regression and growth in tumors. Journal of Theoretical Biology. 2006;241(4):903–918.

3. Dewhirst MW, Cao Y, Moeller B. Cycling hypoxia and free radicals regulate angiogenesis and radiotherapy response. Nature Reviews Cancer. 2008;8(6):425.

4. Stone J, Itin A, Alon T, Pe’Er J, Gnessin H, Chan-Ling T, et al. Development of retinal vasculature is mediated by hypoxia-induced vascular endothelial growth factor (VEGF) expression by neuroglia. Journal of Neuroscience. 1995;15(7):4738–4747.

5. Kerbel RS. Tumor angiogenesis. New England Journal of Medicine. 2008;358(19):2039–2049.

6. Knighton D, Silver I, Hunt T. Regulation of wound-healing angiogenesis-effect of oxygen gradients and inspired oxygen concentration. Surgery. 1981;90(2):262–270.

7. Beer TW, Baldwin HC, Goddard JR, Gallagher PJ, Wright DH. Angiogenesis in pathological and surgical scars. Human pathology. 1998;29(11):1273–1278.

8. Lecoq J, Parpaleix A, Roussakis E, Ducros M, Houssen YG, Vinogradov SA, et al. Simultaneous two-photon imaging of oxygen and blood flow in deep cerebral vessels. Nature medicine. 2011;17(7):893.

9. Krogh A. The number and distribution of capillaries in muscles with calculations of the oxygen pressure head necessary for supplying the tissue. The Journal of physiology. 1919;52(6):409–415.

10. Popel AS. Theory of oxygen transport to tissue. Critical reviews in biomedical engineering. 1989;17(3):257.

11. Lücker A, Weber B, Jenny P. A dynamic model of oxygen transport from capillaries to tissue with moving red blood cells. American Journal of Physiology-Heart and Circulatory Physiology. 2014;.

12. Secomb TW, Hsu R, Dewhirst M, Klitzman B, Gross J. Analysis of oxygen transport to tumor tissue by microvascular networks. International Journal of Radiation Oncology* Biology* Physics. 1993;25(3):481–489.

13. Goldman D, Popel AS. A computational study of the effect of capillary network anastomoses and tortuosity on oxygen transport. Journal of Theoretical Biology. 2000;206(2):181–194.

14. Goldman D, Popel AS. A computational study of the effect of vasomotion on oxygen transport from capillary networks. Journal of Theoretical Biology. 2001;209(2):189–199.

15. Zhu TC, Liu B, Penjweini R. Study of tissue oxygen supply rate in a macroscopic photodynamic therapy singlet oxygen model. Journal of biomedical optics. 2015;20(3):038001.

16. Liu D, Wood N, Witt N, Hughes A, Thom S, Xu X. Computational analysis of oxygen transport in the retinal arterial network. Current eye research. 2009;34(11):945–956.

17. Secomb TW, Hsu R, Park EY, Dewhirst MW. Green’s function methods for analysis of oxygen delivery to tissue by microvascular networks. Annals of biomedical engineering. 2004;32(11):1519–1529.

18. Kreuzer F. Oxygen supply to tissues: the Krogh model and its assumptions. Experientia. 1982;38(12):1415–1426.

19. D’Angelo C. Multiscale modelling of metabolism and transport phenomena in living tissues. EPFL; 2007.

20. Goldman D. Theoretical models of microvascular oxygen transport to tissue. Microcirculation. 2008;15(8):795–811.

21. Cassot F, Lauwers F, Fouard C, Prohaska S, Lauwers-Cances V. A novel three-dimensional computer-assisted method for a quantitative study of microvascular networks of the human cerebral cortex. Microcirculation. 2006;13(1):1–18.

22. Causin P, Guidoboni G, Malgaroli F, Sacco R, Harris A. Blood flow mechanics and oxygen transport and delivery in the retinal microcirculation: multiscale mathematical modeling and numerical simulation. Biomechanics and modeling in mechanobiology. 2016;15(3):525–542.

23. Grimes DR, Kannan P, Warren DR, Markelc B, Bates R, Muschel R, et al. Estimating oxygen distribution from vasculature in three-dimensional tumour tissue. Journal of The Royal Society Interface. 2016;13(116):20160070.

24. Hsu R, Secomb TW. A Green’s function method for analysis of oxygen delivery to tissue by microvascular networks. Mathematical biosciences. 1989;96(1):61–78.

25. Pozrikidis C, Farrow D. A model of fluid flow in solid tumors. Annals of biomedical engineering. 2003;31(2):181–194.

26. Peskin CS. The immersed boundary method. Acta numerica. 2002;11:479–517.

27. Birol G, Wang S, Budzynski E, Wangsa-Wirawan ND, Linsenmeier RA. Oxygen distribution and consumption in the macaque retina. American Journal of Physiology-Heart and Circulatory Physiology. 2007;.

28. Hardarson SH, Harris A, Karlsson RA, Halldorsson GH, Kagemann L, Rechtman E, et al. Automatic retinal oximetry. Investigative ophthalmology & visual science. 2006;47(11):5011–5016.

29. Williamson TH, Rumley A, Lowe G. Blood viscosity, coagulation, and activated protein C resistance in central retinal vein occlusion: a population controlled study. British Journal of Ophthalmology. 1996;80(3):203–208.

30. Riva CE, Grunwald JE, Sinclair SH, Petrig B. Blood velocity and volumetric flow rate in human retinal vessels. Investigative ophthalmology & visual science. 1985;26(8):1124–1132.

31. Funk R. Blood supply of the retina. Ophthalmic research. 1997;29(5):320–325.

